# Epigenetic landscape of the H3K27me3 mark in macrophages transformed by *Theileria annulata*

**DOI:** 10.1101/2025.01.06.631474

**Authors:** Takaya Sakura, Shahin Tajeri, Zineb Rchiad, Hifzur R. Ansari, Abhinav Kaushik, Tobias Mourier, Arnab Pain, Michel Wassef, Gordon Langsley

## Abstract

*Theileria annulata* and *T. parva* are obligate intracellular parasites that induce bovine leukocyte transformation leading to uncontrolled proliferation and heightened dissemination of infected leukocytes. Early passage *T. annulata*-transformed macrophages are virulent, but with long-term culture, they become attenuated for dissemination. Loss of dissemination is restored when TGF-β2-mediated signaling is re-established in attenuated macrophages, suggesting that epigenetic changes might be occurring that contribute to the attenuated phenotype. We focused on one of the important repressive histone marks, tri-methylated lysine 27 of histone H3 (H3K27me3) catalysed by the Polycomb Repressive Complex 2 and found that the global level of H3K27me3 is increased in attenuated macrophages. ChIP-seq revealed the genomic distribution of H3K27me3 is heavily remodeled from virulent to attenuated macrophages, where many genes transition from a focal peak of H3K27me3 around the transcription start site to larger chromatin domains. This transition is not paralleled by large-scale transcriptional downregulation. RNA-seq analysis following PRC2 inhibitor treatment reveals that fewer genes are derepressed in attenuated macrophages than in virulent macrophages, suggesting that broader H3K27me3 profiles do not systematically translate into increased gene silencing activity. Our findings shed light on the mechanisms underlying the dysregulation of epigenetic modifications in *Theileria*-induced leukocyte transformation.

**Significance statement:** Tropical theileriosis is a major constraint to livestock production worldwide. Infection with *Theileria annulata* induces the transformation of bovine leukocytes, which subsequently exhibit similarities to human leukemia. Long-term *in vitro* culture of transformed leukocytes attenuates their tumorigenic properties. The molecular changes underlying this process are currently unclear. We focused on H3K27me3, a key histone tail post-translational modification that regulates chromatin structure and gene expression, and compared its genomic landscapes in virulent, highly disseminating macrophages *versus* attenuated macrophages with reduced dissemination potential that are used as a live vaccine against tropical theileriosis. Western blot and ChIP-seq revealed higher global levels of H3K27me3 and a striking broadening of its genomic distribution in attenuated macrophages. To our surprise, this reconfiguration is not associated with major changes in gene expression and attenuated cells appear to be less responsive to PRC2 inhibition, suggesting that large-scale broadening of H3K27me3 does not systematically translate into increased PRC2-mediated silencing activity. Despite these changes, H3K27me3 maintains the silencing of key tumor suppressor genes in both virulent and attenuated macrophages. These findings shed light on the interplay between infection, the evolution of oncogenic potential and reconfiguration of the chromatin landscape following attenuation of virulence.

## Introduction

Apicomplexan parasites *Theileria annulata* and *T. parva* are transmitted by ticks and infect bovine leukocytes, causing tropical theileriosis primarily in Asian countries and East Coast Fever (ECF) in Africa^1,2^. Macroschizonts differentiated from tick-injected sporozoites induce tumorigenesis of infected leukocytes, and dissemination of transformed leukocytes is mainly responsible for the pathology of these livestock diseases. Despite the huge economic impact of this disease in the veterinary field, available vaccines suffer from several drawbacks such as the requirement for quality control and cold chain^3,4^. In addition, parasites resistant to the theileriacidal drug buparvaquone are emerging in endemic areas^5,6^. New vaccines and drugs are therefore needed to control these widespread livestock diseases and to reduce their economic burden in endemic areas. A specific feature of *T. annulata* and *T. parva* infections is the transformation of their host leukocytes that allows the parasite population to grow due to uncontrolled host cell proliferation^7,8^. The mechanisms underpinning *Theileria annulata*-mediated transformation – i.e., how parasites modulate host gene expression and hijack their proliferation – rely on host-parasite interactions. For instance, the IκB kinase (IKK) complex is recruited to the surface of the parasite resulting in nuclear factor kappa B (NF-κB) being constitutively activated to induce expression of anti-apoptotic genes^9^. In addition, *Theileria* infection inhibits nuclear translocation of p53, a well-known tumor suppressor, and blocks p53-driven expression of pro-apoptotic genes^10,11^. Meanwhile, c-Jun N-terminal Kinase (JNK2) interacts with a GPI-anchored macroschizont surface protein (p104), while JNK1 translocates to the nucleus to activate c-Jun to drive transcription of the matrix metalloproteinase 9 (*mmp*9) gene and enhance tumor dissemination^12–14^. Parasite secreted peptidyl-prolyl isomerase PIN1 also contributes to c-Jun stabilization that maintains the proliferative phenotype of *Theileria*-infected leukocytes^15^. Subversion of host cell signal transduction pathways augments *c-myc*-driven transcription of anti-apoptotic genes^16–18^. *Theileria* transformation therefore shares several similarities with human carcinogenesis, where epigenetic (i.e., not involving changes of the DNA sequence) alterations contribute to cancer development. For example, *Theileria*-induced tumor dissemination is regulated by changes in TGF-β2 levels and dissemination of attenuated macrophages is restored by addition of exogenous TGF-β2 consistent with a role for epigenetic regulation of cancer-related genes^19–21^.

Many types of post-translational modifications on histone tails have been shown to regulate chromatin dynamics and transcription^22^. Changes of the chromatin state have been reported in *Theileria-*infected leukocytes. Tri-methylated lysine 4 of histone H3 (H3K4me3) is a mark correlating with active transcription enriched upstream of the *mmp-9* gene in the *Theileria*-infected B-cell lymphosarcoma cell line (TBL3), and transcriptional activation of *mmp-9* is associated with heightened tumor dissemination^23,24^. Tri-methylated lysine 27 of histone H3 at lysine 27 (H3K27me3) is a key chromatin modification catalyzed by the Polycomb Repressive Complex 2 (PRC2)^25^. H3K27me3 is associated with transcriptional silencing, acting in part as a docking site for CBX-containing PRC1 complexes. During embryonic development and throughout adult life, H3K27me3 plays key a role in maintaining transcriptional silencing of genes encoding cell fate regulators^26^. Furthermore, the PRC2 complex is altered in several types of tumors, acting either as a tumor suppressor or an oncogene^27^.

*Theileria annulata*-infected macrophages are fully transformed and as such can be maintained in culture indefinitely. Early passage *T. annulata*-transformed macrophages are virulent (disseminating), but virulence decreases with long-term (>300) passage and infected cells become attenuated for dissemination^28,29^. In the present study, using the Polycomb-specific H3K27me3 histone mark as a paradigm, we investigate the evolution of the epigenetic landscape between virulent and attenuated *T. annulata*-infected macrophages. We show that the distribution of H3K27me3 in attenuated macrophages is profoundly altered and that attenuated cells become less responsive to PRC2 inhibition compared to virulent macrophages. However, some tumor suppressor genes still respond to PRC2 inhibitor treatment. Our study therefore reveals profound changes in the chromatin landscape that accompany the transition from virulent to attenuated states.

## Results

### *Theileria annulata* parasites within transformed host leukocytes are not stained by a H3K27me3-specific antibody

When comparing the amino acid sequence of the tail of histone H3 between various parasite species, we found that the amino acid sequence surrounding Lys27 is highly conserved between mammals and *Plasmodium falciparum* and *Toxoplasma gondii* (Fig. 1a). Accordingly, H3K27me3 is present in *T. gondii* tachyzoites and has been detected in sexual stage II gametocytes in *P. falciparum*^30–33^. We noticed that there is an S>T change at position 28 of *Theileria* parasite histone H3, raising the question whether H3K27 is tri-methylated in *Theileria* species. As expected, immunofluorescence analysis using an antibody specific for H3K27me3 stained bovine host nuclei, whereas *Theileria* nuclei were not stained (Fig. 1b). Whether this is due to absence of the mark and/or a failure of the antibody to recognize it (due to the H3S28>T change) is unclear. By contrast, specific antibodies to H3K4me3 stained both parasite and host cell nuclei. Since H3K27me3 is not detected in the parasite by the antibody we used, the following analyses of H3K27me3 reflect only the bovine histone mark.

**Figure 1.**
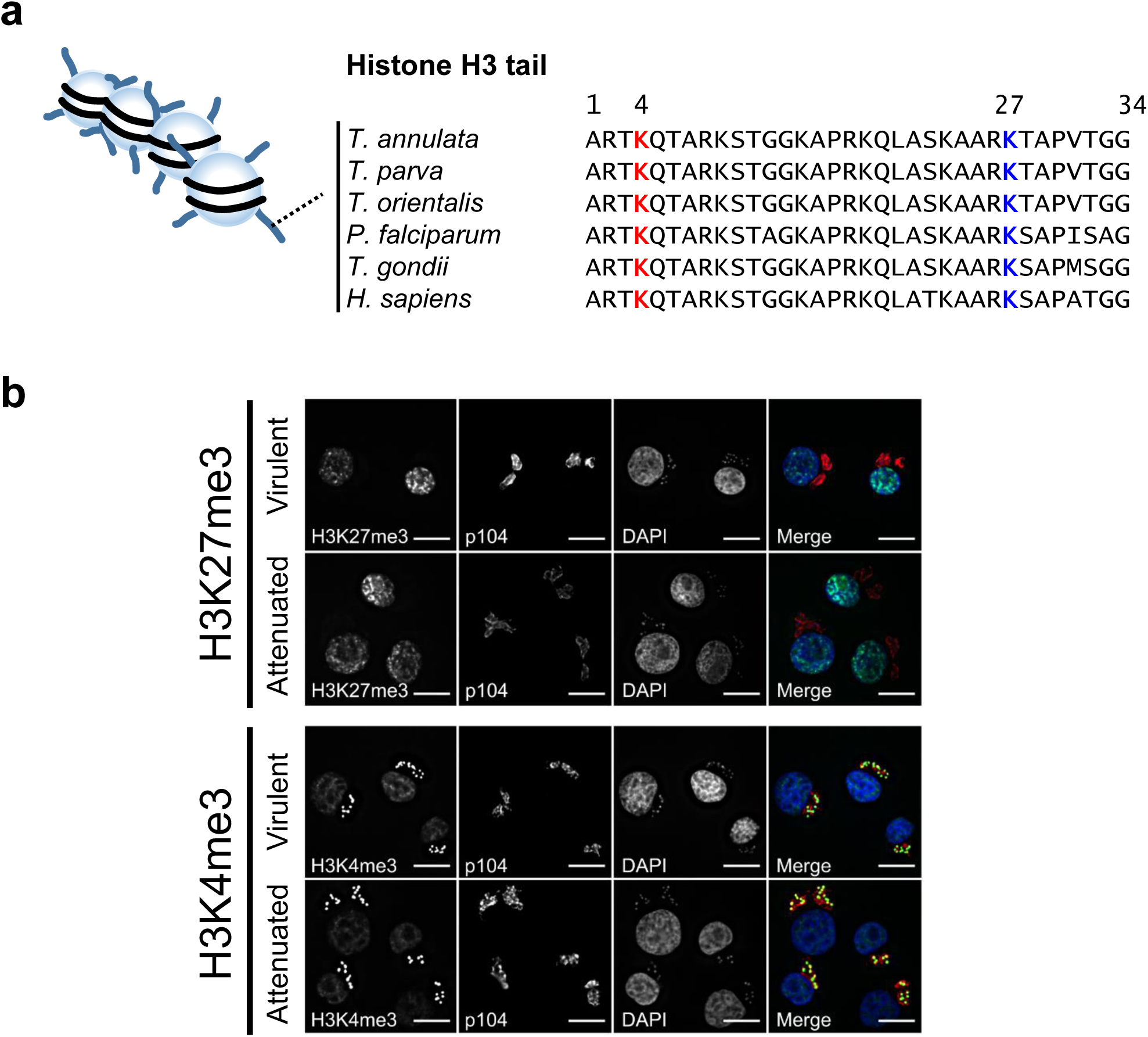
Localization of H3K27me3 and H3K4me3 marks in *Theileria*-infected virulent and attenuated macrophages. **a)** Amino acid sequence alignment of Histone H3 tail among Apicomplexan parasites and *Homo sapiens*. TA07840 sequence is used for *Theileria annulata.* **b)** Immunofluorescence staining with H3K27me3 and H3K4me3 (green). The monoclonal p104 antibody labels a parasite surface protein p104 (red) and DAPI stained nuclei (blue). H3K27me3 antibody only stains host nuclei. Tri-methylation on lysin 4 is conserved between *T. annulata* and bovine host nuclei. Scale bar: 10 μm

### Global levels and distribution of H3K27me3 are altered in attenuated macrophages

The fact that attenuated macrophages have a low dissemination potential led us to hypothesize that oncogenes activated as a consequence of *Theileria* infection might be transcriptionally downregulated through mechanisms promoting a repressed chromatin state. To this end, we quantified H3K27me3 abundance in the nuclear fraction of both virulent and attenuated macrophages (Fig. 2a). The amount of H3K27me3 was estimated to be 1.58-fold higher in attenuated macrophages compared to virulent macrophages, whereas H3K4me3 was 0.66-fold lower in attenuated macrophages. Interestingly, the expression of EZH2, the main PRC2 catalytic subunit, appeared unchanged between virulent and attenuated macrophages.

**Figure 2.**
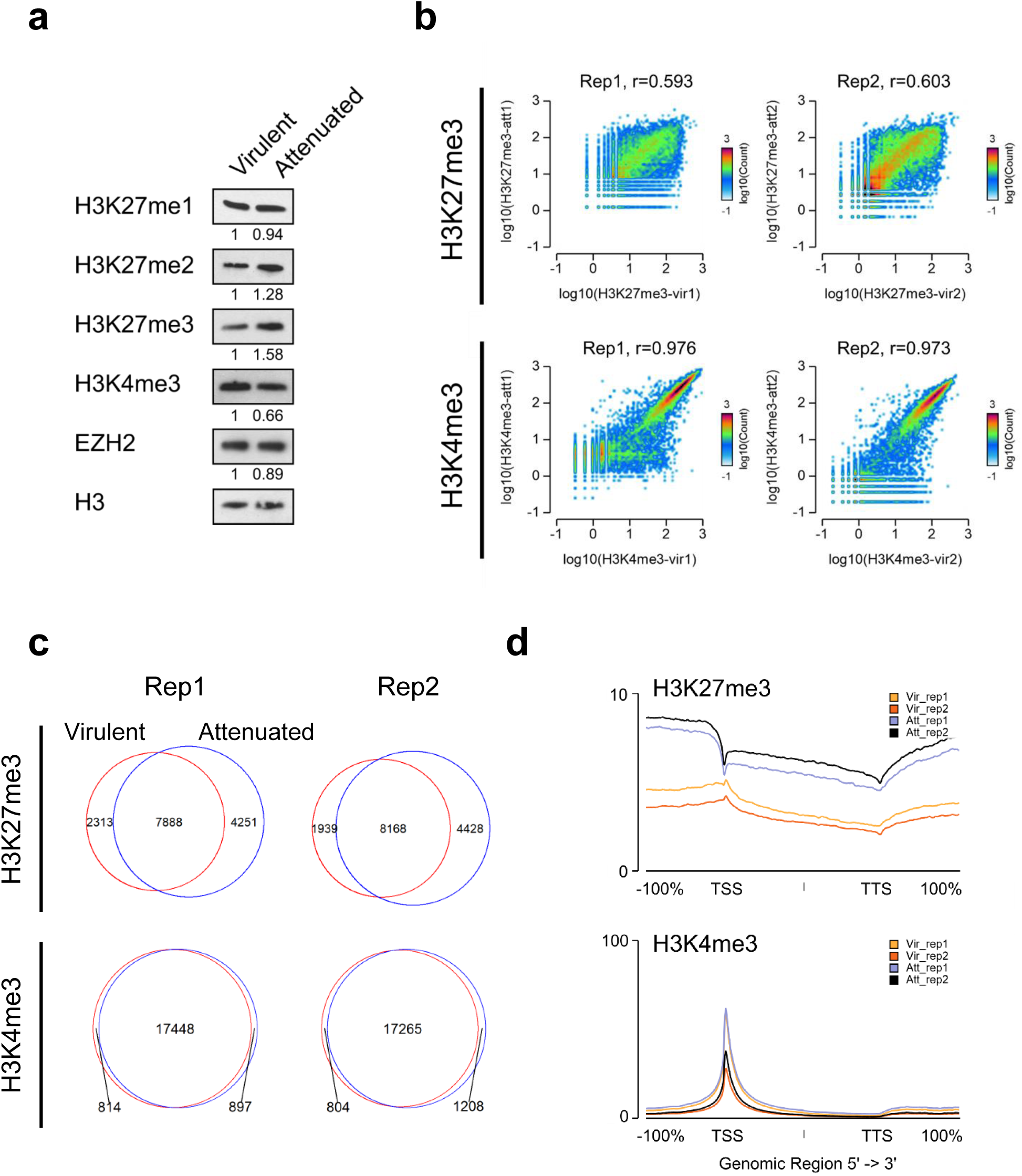
Genome-wide H3K27me3 distribution dramatically changes in attenuated macrophages. **a)** Comparative abundance of histone marks and EZH2 in virulent and attenuated macrophages. Each band was normalized to H3, used as a loading control. Relative intensity is shown below each band. **b)** 2D density plots comparing H3K27me3 and H3K4me3 signal between virulent and attenuated macrophages in each of the two biological replicates. Read counts for each gene in virulent (X-axis) and attenuated (on Y-axis) macrophages are plotted. The r values represent the Pearson correlation coefficients. **c)** Venn diagrams showing the overlap between virulent and attenuated macrophages for H3K27me3- and H3K4me3-positive genes. Peaks of H3K27me3 and H3K4me3 were associated to the *B. taurus* transcripts (UMD3.1.1) and signal intensity around the TSS +/- 2 kb was quantified using Easeq (52). **d)** Average H3K27me3 and H3K4me3 signal intensities at all annotated *Bos taurus* genes (bosTau8). The X-axis represents a window centered on each gene, spanning from the transcription start site (TSS) to the transcription termination site (TTS) with an additional ±100% flanking region. Yellow/orange lines represent signals from virulent macrophages, whereas purple/black lines represent those from attenuated macrophages.

We next investigated the genome-wide distribution of the two histone marks in *Theileria*-infected macrophages by chromatin immunoprecipitation sequencing (ChIP-seq). *Drosophila melanogaster* chromatin was used as a spike-in to normalize the abundance of the histone marks^34^. Consistent with immunoblotting results, the density of H3K27me3 around transcriptional start sites (TSS) was generally higher in attenuated than virulent macrophages, while H3K4me3 density was lower (Supplementary Fig. 1a). H3K27me3 was also poorly correlated between virulent and attenuated macrophages at the genome-wide level (r=0.593 and 0.603 for replicates 1 and 2, respectively), contrasting with the high degree of H3K4me3 correlation (r=0.976 and 0.973 for replicate 1 and 2 respectively) (Fig. 2b). A substantial fraction of H3K27me3-positive genes were specific to either virulent or attenuated macrophages, while a relatively smaller proportion of H3K4me3-positive genes differed between the two cell states (Fig. 2c). Strikingly, plotting the average H3K27me3 signal revealed that while the mark was increased overall over the gene bodies of attenuated cells, its relative distribution around the TSS was markedly different between virulent and attenuated cells (Fig. 2d). In virulent macrophages, H3K27me3 showed a typical enrichment around the TSS. However, in attenuated macrophages, H3K27me3 enrichment was no longer maximal at the TSS. Instead, the mark accumulated most prominently in intergenic regions (upstream of TSS and downstream of transcription termination sites). This contrasts with H3K4me3 that accumulated sharply around the TSS and showed a similar distribution pattern between virulent and attenuated macrophages.

We further defined 3 clusters of genes according to H3K27me3 enrichment (Fig. 3a): virulent-positive (cluster 1), attenuated-positive (cluster 2), both positive (cluster 3). While virulent-specific H3K27me3 signal showed a sharp enrichment around the TSS, attenuated-specific H3K27me3 signal showed a wide distribution around the TSS, with a maximal enrichment upstream of the TSS. This difference was also obvious for genes showing H3K27me3 enrichment in both conditions (Fig. 3a). Average transcript levels in virulent and attenuated macrophages for each H3K27me3 cluster showed no statistically significant difference as a group, while differential gene expression analysis identified a subset of cluster 2 genes that were downregulated in attenuated *versus* virulent macrophages (Fig. 3b)^35^. One of the genes highly enriched for H3K27me3 in cluster 2 of attenuated macrophages is the Src Kinase Associated phosphoprotein 2 (*SKAP2*), which is associated with macrophage cell adhesion ^36^. It is covered by a large domain of H3K27me3 extending over 500 kb, in contrast to a low level concentrated at the TSS found in virulent macrophages (Fig. 3c). These differences are paralleled by a corresponding downregulation of *SKAP2* expression in attenuated macrophages^35^. Patterns of H3K27me3 and H3K4me3 distribution around representative genes of other clusters, such as TBX5 (Cluster 1) and WNT3 (Cluster 3), further illustrate the broader spreading of H3K27me3 in attenuated cells compared to virulent cells (Fig. 3c). Altogether, these data show that the attenuated phenotype is characterized by a dramatic reconfiguration of H3K27me3 distribution.

**Figure 3.**
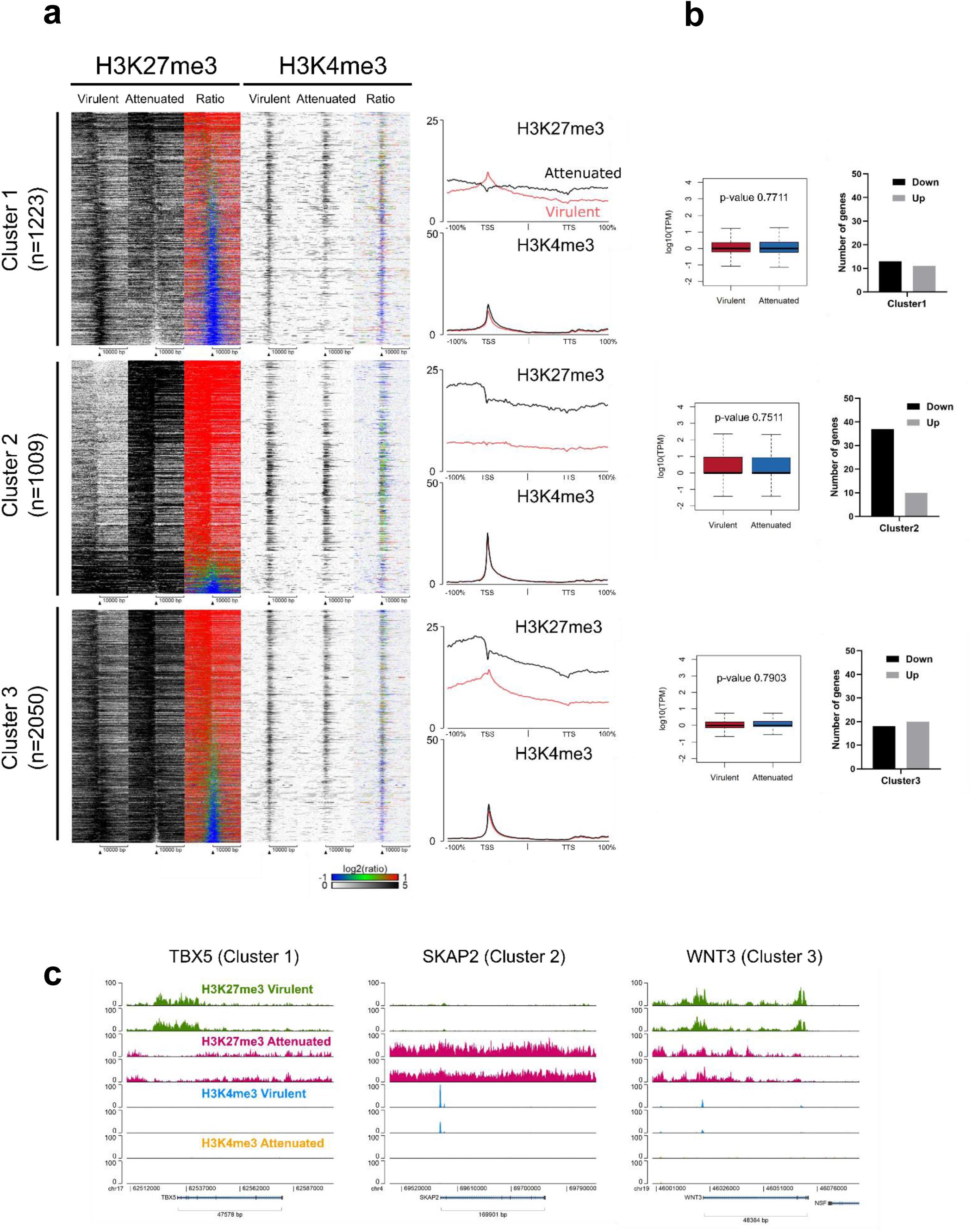
Association between shifts of H3K27me3 profile and gene expression. **a)** H3K27me3 and H3K4me3 heat maps around TSS +/- 10 kb of H3K27me3-positive genes either in virulent or attenuated macrophages. Genes were sorted by the log2-transformed ratio of H3K27me3 signal between attenuated and virulent macrophages (Att/Vir). Cluster 1: positive in virulent macrophages, Cluster 2: positive in attenuated macrophages, Cluster 3: positive in both macrophages. Replicate 2 heat maps are shown. Average H3K27me3 and H3K4me3 signal intensities of the genes of the corresponding clusters are shown on the right of each heat map. The X-axis represents a window centered on each gene, spanning from the transcription start site (TSS) to the transcription termination site (TTS) with an additional ±100% flanking region. Black lines represent signals from virulent macrophages, whereas red lines represent those from attenuated macrophages. **b)** Gene expression levels (box plots, left) and DEGs (histograms, right) of each cluster. Average TPM values of each cluster were compared between virulent and attenuated macrophages and p-values were calculated using Mann-Whitney U test. Within each cluster, the number of down or up DEGs in attenuated *versus* virulent macrophages was determined using DESeq2 (FDR < 0.1, p-value < 0.05, logFC > 1, logFC < −1). **c)** Genomic distribution of H3K27me3 and H3K4me3 around representative genes of each cluster.

### Limited effect of PRC2 inhibition on tumor-related phenotypes

In light of the widespread re-distribution of H3K27me3 in attenuated cells, we next investigated the requirement for the mark in the transformed and attenuated phenotypes. We treated both virulent and attenuated macrophages with a specific PRC2 inhibitor (UNC1999) or its inactive analog (UNC2400)^37,38^. Ten days treatment with UNC1999 led to an acute loss of H3K27me3 in both cell lines (Fig. 4a). We then performed fibronectin-binding assays on the PRC2 inhibitor treated macrophages^19^. Though attenuated macrophages displayed lower binding activity compared with virulent macrophages, PRC2 inhibition did not affect binding activity (Fig. 4b). We injected the PRC2 inhibitor treated macrophages into immune-compromised Rag2/γC mice and then orally administered PRC2 inhibitor for 3 weeks to evaluate if loss of H3K27me3 impacts the formation of tumors *in vivo*. The parasite burden of heart and lung was quantified by measuring the amount of DNA corresponding to the *T. annulata ama-1* gene (TA02980). Similar to the fibronectin-binding assay, there was no significant change resulting from PRC2 inhibition (Fig. 4c).

**Figure 4.**
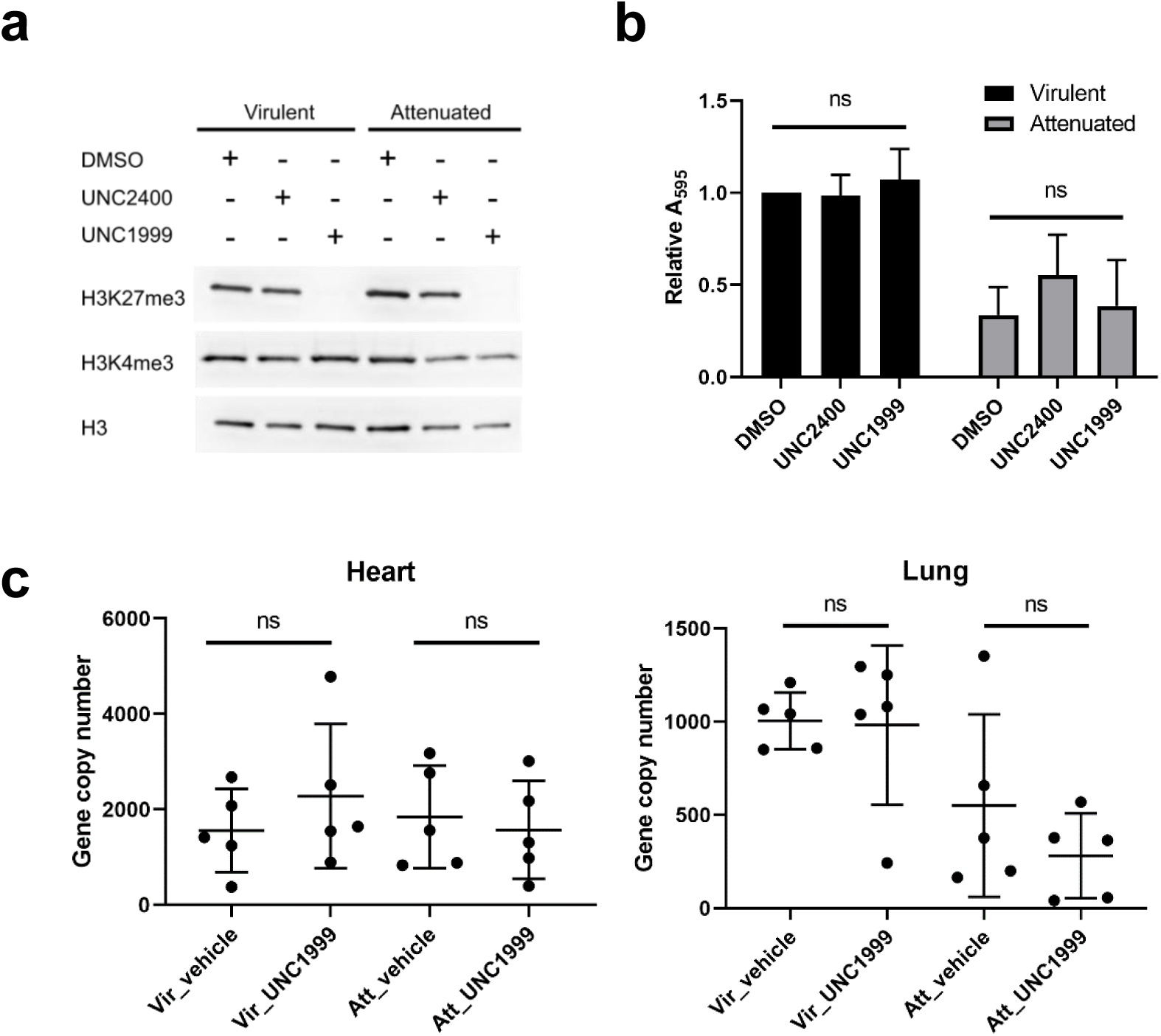
PRC2 inhibition does not alter tumour dissemination properties of virulent or attenuated cells. **a)** PRC2 inhibitor treatment (UNC1999) lead to loss of the H3K27me3 histone mark, whereas the inactive analog UNC2400 had no detectable effect on H3K27me3 levels. Both macrophages were treated with DMSO or PRC2 inhibitors for 10 days and nuclear extracts were prepared. **b)** Fibronectin binding assay. Cells treated with PRC2 inhibitor were incubated in a fibronectin coated well plate for 1 hour at 37 °C. Error bars show ± SD from 3 biological replicates. **c)** Dissemination of *T. annulata*-infected macrophages in immune-compromised Rag2/γC mice. Gene copy number of parasites in each organ were obtained by absolute quantification of the *T. annulata ama-1* gene^80^.

### H3K27me3-mediated gene repression is dampened in attenuated macrophages

We next assessed gene expression changes following PRC2 inhibition in virulent and attenuated cells. For this purpose, we performed RNA-seq on *Theileria*-infected macrophages treated with UNC1999 or UNC2400 for 10 days to identify genes whose transcription is reactivated following loss of PRC2 activity. Consistent with the repressive function of H3K27me3, PRC2 inhibition resulted in a majority of differentially expressed genes (DEGs) being upregulated in both virulent and attenuated macrophages (Fig. 5a). Interestingly, the number of upregulated genes in virulent macrophages was more than 2-fold higher than in attenuated macrophages, suggesting that gene silencing is less dependent on PRC2 in attenuated macrophages. Twenty-one genes were commonly upregulated in virulent and attenuated macrophages, suggestive of a core function for H3K27me3-mediated repression in *Theileria*-infected macrophages (Fig. 5b, Supplementary Fig. 2, 3, and Supplementary Table 2). Gene ontology analysis also indicated that PRC2 retains its canonical function of regulating genes related to embryonic development in virulent macrophages. However, enrichment for this class of genes is no longer statistically significant in attenuated macrophages (Fig. 5c and Supplementary Table 4). This functional difference in PRC2 inhibitor responsiveness between virulent and attenuated macrophages could be caused by the difference of genome-wide distribution of H3K27me3 (Fig. 2b). Remarkably, some of the common PRC2-repressed genes (Fig. 5d) include Granzyme A (*GZMA*) identified as a novel dissemination suppressor of *Theileria*-transformed macrophages^35^ and Follistatin (*FST*) known to reduce tumor invasiveness and metastasis in breast cancer^39,40^ (Supplementary Fig. 2).

**Figure 5.**
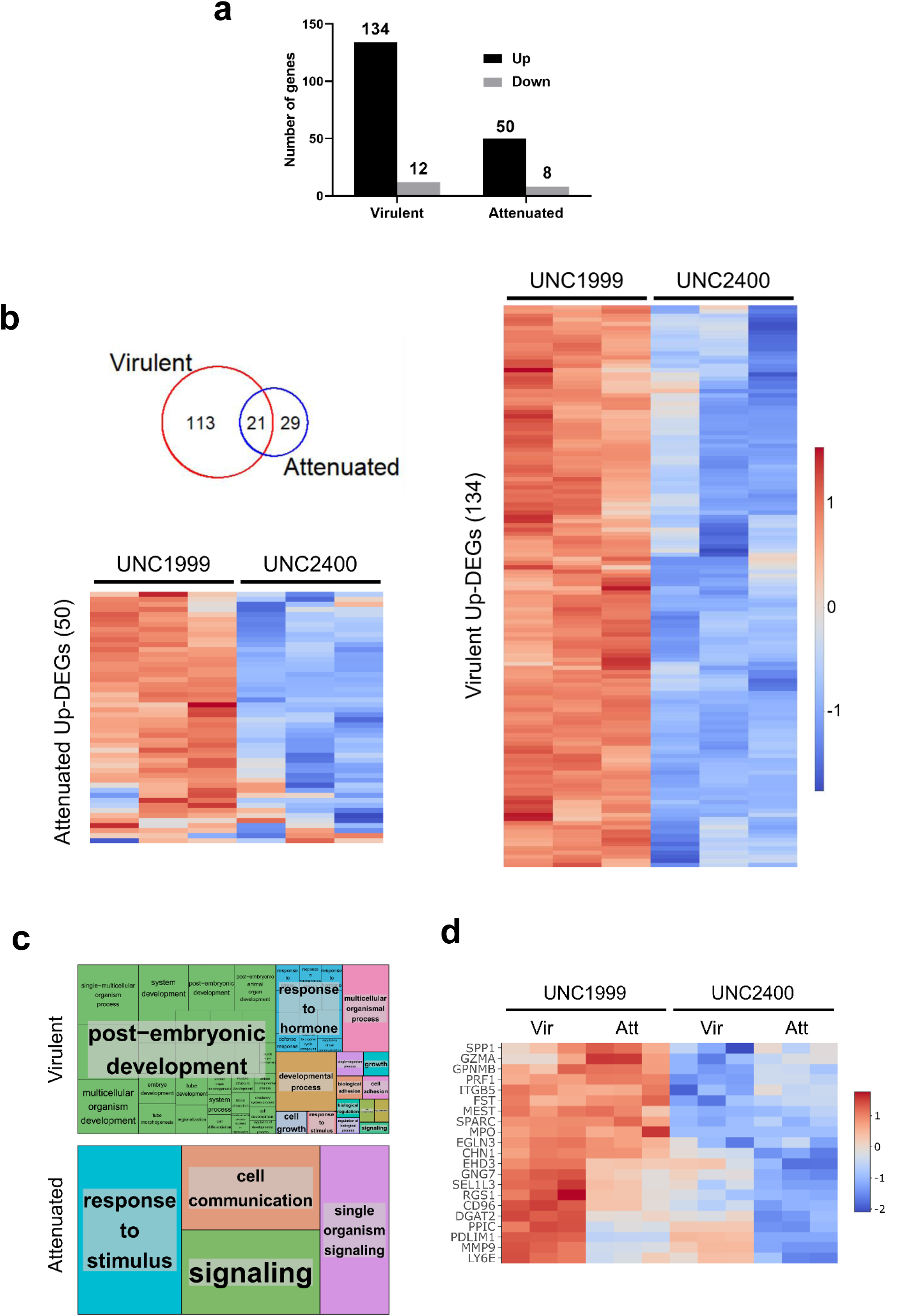
H3K27me3-mediated gene repression is dampened in attenuated macrophages. **a)** Number of DEGs (logFC > 1, logFC < −1, padj < 0.05) between virulent and attenuated macrophages following PRC2 inhibitor treatment. **b)** Venn diagram of common and uniquely upregulated genes in both virulent (134 genes) and attenuated (50 genes) macrophages with the heatmaps of upregulated DEGs. **c)** Gene ontology analysis. Treemap were visualized by Revigo^85^. These pathways were specifically upregulated upon PRC2 inhibitor treatment in each macrophage. **d)** Heat map of the 21 common upregulated genes in both macrophages upon UNC1999 treatment versus inactive analog UNC2400.

Overall, our analyses of the epigenetic landscape of *Theileria*-infected macrophages have uncovered a dramatic reconfiguration of the H3K27me3 landscape during the transition from the virulent to the attenuated state. The histone mark adopts a broad pattern of genomic distribution reminiscent of the transition from sharp to broad domains observed during development in the transition from the pluripotent to the differentiated state^41^. This is paralleled by a reduced transcriptional response following PRC2 inhibition, suggesting that this broad pattern may be less conducive to gene silencing and/or more resilient to PRC2 inhibition. Interestingly, silencing of some key tumor suppressor genes is preserved in attenuated macrophages, highlighting the requirement for a minimally functional PRC2 complex.

## Discussion

The unrestrained leukocyte proliferation and illicit dissemination within *Theileria*-infected animals have several similarities with human carcinogenesis, and for this reason we focused on key post-translational modifications of histone tails with established links to oncogenic phenotypes. We chose to map H3K4me3 and H3K27me3 histone marks in virulent disseminating *T. annulata*-transformed versus attenuated macrophages with dampened dissemination^42^. As H3K27me3 is a key repressive mark we hypothesized that changes in its genomic distribution could be involved in the attenuation processes. Indeed, we observed that the distribution of H3K27me3 is dramatically reconfigured during attenuation. In virulent macrophages, H3K27me3 preferentially accumulates around gene TSSs, while in attenuated macrophages the histone mark adopts a broader profile over large regions sometimes encompassing multiple genes (Fig. 2d and Fig. 3a/c). Furthermore, H3K27me3 peaks on the DEGs between virulent and attenuated macrophages also exhibited a similar distribution (Supplementary Fig. 4)^35^.

Given that follicular lymphoma harboring EZH2 mutations also display a broadened H3K27me3 methylation profile, we looked at SNPs in both *ezh1* and *ezh2* genes of virulent and attenuated macrophages (data not shown), but could find no evidence of any amino acid changes. Long-term oxidative stress can also lead to increases of H3K27me3 levels and attenuated macrophages exhibit elevated H_2_O_2_ production^43,44^. In fact, a number of different possible mechanisms could be involved, reviewed in^45,46^. For example, expression of PRC2 subunits and co-factors can increase relative to cell proliferation rate, but our RNA-seq data indicate that any differences are very subtle, as are changes in expression of H3K27me3 demethylases KDM6A (very slightly down) and KDM6B (very slightly up) in attenuated macrophages. Increased abundance in attenuated macrophages of H3K27ac or H3K36me3 marks might directly increase PRC2 activity, and decreased DNA methylation can also lead to an increase in PRC1 activity. Additionally, *T. annulata* is known to secrete a number of proteins predicted to be <u>h</u>ighly intrinsically <u>d</u>isordered <u>p</u>roteins (HIDP) such as TashAT2^47^, Ta9^48^ and the NIDP family^49^, and all have been located to the host cell nucleus. Interestingly, in the context of chromatin silencing, overexpression of TashAT2 in BoMacs repressed transcription of 518 genes and upregulated transcription of 319 genes out of 837 DEGs^47^. Moreover, when Ta9 was overexpressed in BoMacs^50^ the top-scoring transcription factor predicted to regulate Ta9-responsive DEGs was MTF2, a component of the PRC2 complex^51^. So, secreted parasite HIDPs might contribute to the formation of phase-separated compartments in the chromatin of host cells^52,53^.

While these represent potential drivers of H3K27me3 spreading in attenuated macrophages, such spreading may arise from a combination of mechanisms.

Interestingly and importantly, we found that reconfiguration of H3K27me3 distribution is not associated with widespread gene silencing and that fewer genes become de-repressed following PRC2 inhibition in attenuated cells than in virulent macrophages. The 50 DEGs identified in attenuated macrophages through RNA-seq following PRC2 inhibitor treatment were integrated with ChIP-seq. Of these, eight genes (CD69, IRF1, TMEM108, ENC1, CHN1, EHD3, GZMA, LY6E) were classified within cluster 2 in Fig. 3a, while the remaining genes were categorized into clusters 1 and 3. However, these analyses did not reveal any distinct chromatin features (Supplementary Fig. 3). These observations suggest that reconfiguration of H3K27me3 does not lead to a globally increased PRC2 silencing activity. Why spreading of H3K27me3 does not lead to a more extensive repression of transcription in attenuated macrophages is unclear, but we note that genes that gain H3K27me3 in attenuated macrophages are on average already lowly expressed in virulent cells (Fig. 3b). Previous studies have shown that global increases in H3K27me3 can lead to complex effects on transcription that do not necessarily result in a stronger repression of Polycomb target genes^54,55^. Interestingly however, PRC2 inhibitor treatment led to the de-repression of three tumor suppressor genes *GZMA, PRF1* and *FST* in both virulent and attenuated macrophages, indicating that H3K27me3 is required to maintain their repressed state, thus potentially contributing to the cancer-like phenotype of *Theileria-*induced transformation. This is consistent with our observation that *GZMA* plays a role in tumor dissemination of *Theileria*-infected leukocytes and human B-cell lymphomas^35^. The observation that genes silenced by PRC2 comprise both tumor suppressors and oncogenes may explain the lack of any obvious effect of inhibiting PRC2 on the transformed phenotype. Alterations of the chromatin state following host-parasite infection have also been reported in other contexts. In particular, intracellular pathogens are capable of epigenetically modifying gene expression of infected host cells by injecting effector molecules. *Legionella pneumophila* secretes the methyltransferase RomA into the host cell to modify H3K14 acetylation and promote intracellular bacterial replication^56^. Amastigotes of *Leishmania*, an obligate intracellular parasite, can induce epigenetic alterations of host macrophages, such as DNA and histone modifications to promote pathogen survival and replication^57^. For instance, modulation of host H3 acetylation and methylation by *Leishmania* amastigotes prevents NF-kB and NLRP3 inflammasome activation, resulting in suppression of host immune responses^58,59^. It is noteworthy that *Toxoplasma* E2F4-associated EZH2 inducing Gene Regulator (TEEGR) activates EZH2 and antagonizes NF-κB signaling by epigenetic repression in host cells^60^. Although host EZH2 protein levels appeared equivalent in attenuated macrophages (Fig. 2a), other mechanisms such as phosphorylation^61,62^ and oxidative stress^43,44^ may contribute to heightened methyltransferase activity and/or decreased demethylase activity, thus resulting in spreading of the H3K27me3 mark in attenuated macrophages.

Evaluating H3K27me3 levels in non-infected leukocytes would be informative. However, uninfected primary macrophages do not proliferate unless stimulated and therefore would be difficult to use as a negative control for the analysis of H3K27me3 levels and genomic distribution. In fact, having a good negative uninfected control is a problem when studying *Theileria*-induced macrophage transformation, as more generally it is for cancer research, where the primary tumour progenitor is often not available. For *Theileria*-transformed macrophages BoMacs are occasionally taken as a negative control^63^, but BoMacs have been immortalized by SV40 virus^64^. Thus, taking BoMacs as a negative control would entail comparing the H3K27me3 levels/distribution in SV40-immortalized macrophages with that observed in *Theileria*-transformed macrophages, which is far from ideal. Occasionally in *Theileria* research, uninfected yet immortalized BL3 or BL20 cells are used as negative controls especially when testing parasite-specificity of drug doses^65,66^. We compared H3K27me3 levels between BL3 and *T. annulata*-infected BL3 (TBL3), which revealed that infection is associated to a reduction in H3K27me3 levels (Supplementary Fig. 5, left). However, as our focus was on determining if there was a role for PRC2-mediated silencing of virulence in attenuated Ode macrophages we reasoned that the best control to for attenuated Ode macrophages was to compare the distribution of H3K27me3 marks with that of virulent Ode macrophages, as they are isogenic and harbour the same *T. annulata* parasite strain. To that end, virulent *T. annulata*-transformed macrophages were treated with buparvaquone (BPQ), following the protocol described in^67^, to selectively eliminate the parasite without affecting host cell viability. Total RNA was extracted, and the expression levels of bovine *ezh1* and *ezh2,* as well as parasite p104 (a control for parasite clearance) were assessed by qRT-PCR (Supplementary Fig. 5, right). *Theileria* infection appears to lead to a downregulation of both *ezh1* and *ezh2* and this may be a first step in infection-induced macrophage dedifferentiation^68,69^. Moreover, it might also enable parasite infection to rewire host cell gene expression^70^ and the increase in H3K4me3 signal (at the exposure used) is consistent with rewiring of the host transcriptome.

Our roadmap for future research however, does not include further characterization of PRC2-mediated H3K27me3 silencing, since our *in vivo* mice study suggests that inhibiting PRC2 activity is not a clinically viable control strategy (Fig. 4). PRC2-inhibitor treatment had no effect on *Theileria*-induced tumour dissemination in Rag2gC mice and its application to treat cattle is difficult to imagine given the large quantities required. Rather, we propose to concentrate on other repressive marks that could potentially play a role in the loss of *Theileria*-transformed macrophage virulence, with a particular interest in H3K9me3^71^ for its potential role in repressing the expression of specific host and/or parasite virulence genes^72^.

## Materials and Methods

### Cell culture

*Theileria annulata*-infected macrophages are the Ode virulent and attenuated lines corresponding to passage 53 and 309 ^29,73^. These cells were maintained with RPMI medium supplemented with 10% fetal bovine serum (FBS), 2 mM L-Glutamine, 100 U penicillin, 0.1 mg/ml streptomycin, and 4-(2-hydroxyethyl)-1-piperazineethanesulfonic acid (HEPES) at 37°C with 5% CO_2_.

### Immunofluorescence microscopy

1×10^5^ cells were centrifuged on glass slide with the Cellspin I (Tharmac) at 1,500 rpm for 3 min and fixed by 4% paraformaldehyde for 10–15 min at room temperature. Fixed cells were permeabilized with 0.2% Triton X-100 for 10 min and blocked with 1% BSA for 30 min. These cells were incubated with primary antibodies against H3K27me3 (1/1000, Cell Signaling) or H3K4me3 (1/1000, Cell Signaling) with parasite surface protein p104 (1/1000, 1C12)^74^ for 1 h at room temperature, sequentially stained with secondary antibodies conjugated with Alexa 488 and Alexa 594 (1/1000, Molecular Probes) for 45 min at room temperature. Stained cells were mounted in ProLong Diamond Antifade Mountant with DAPI (Thermo Fisher Scientific). Stained cells were imaged by wide field Leica DMI6000 and these images were deconvoluted.

### Preparation of nuclear extracts

We washed 0.5–2 × 10^7^ *T. annulata*-infected macrophages and *D. melanogaster* S2 cells with phosphate-buffered saline (PBS) once and subsequently suspended with Buffer A (10 mM pH7.9 HEPES, 5 mM MgCl_2_, 250 mM sucrose, 0.1% NP40) supplemented with protease inhibitors and 1 mM DTT. After keeping on ice for 10 min, cell suspensions were centrifuged 8,000 rpm for 10 min at 4°C to collect cytosolic fraction. Pellets were subsequently suspended with Buffer B (25 mM pH7.9 HEPES, 1.5 mM MgCl_2_, 0.1 mM EDTA, 20% Glycerol, 700 mM NaCl) supplemented with protease inhibitors and 1 mM DTT followed by sonication for 10 min on ice. Following centrifugation (14,000 rpm for 15 min at 4 °C), nuclear fractions were collected from the supernatant. Collected nuclear extracts were quantified using the Bradford protein assay.

### Immunoblotting

Nuclear extracts were run by SDS-PAGE and transferred to nitrocellulose membrane (GE healthcare), subsequently membranes were blocked with 5% (w/v) skimmed milk powder in PBS containing 0.1% Tween 20 (PBS-T) for 1 h at room temperature. Immune reactions were carried out using 1/2000 diluted monoclonal rabbit anti-H3K27me3 or H3K4me3 antibodies (Cell signaling) or 1/50000 diluted monoclonal rabbit anti-H3 antibodies (Active motif) overnight at 4 °C as primary antibody. 1/5000 diluted HRP-labeled anti-rabbit secondary antibodies (Santa Cruz) for 1 h at room temperature. Proteins were visualized with X-ray film or ECL (Thermo Fisher Scientific) and imaged fusion FX (Vilber Lourmat). The H3 levels were used as a loading control.

### Preparation of chromatin

We cross-linked 1.5 × 10^7^ cells by pre-warmed DMEM supplemented with 1% Formaldehyde, 15 mM NaCl, 150 μM EDTA, 75 μM EGTA, and 15 mM pH 8.0 Hepes for 10 min at room temperature with slow agitation and then quenched formaldehyde with 125 mM Glycine for 5 min at room temperature. Following centrifugation at 1000 rpm for 10 min at 4 °C, cells were washed with cold PBS once. Washed cells were suspended with 1 ml Buffer 1 (50 mM pH7.5 Hepes-KOH, 140 mM NaCl, 1 mM EDTA, 10% glycerol, 0.5% NP40, 0.25% Triton X-100) supplemented with protease inhibitors (PMSF, aprotinin, leupeptin, pepstatin) and rocked at 4 °C for 10 min. Following centrifugation at 1500 rpm for 5 min at 4 °C, cells were resuspended with 1 ml Buffer 2 (10 mM pH8 Tris, 200 mM NaCl, 1 mM EDTA, 0.5 mM EGTA) supplemented with protease inhibitors (PMSF, aprotinin, leupeptin, pepstatin) and rocked at room temperature for 10 min. Cells were centrifuged at 1500 rpm for 5 min at 4 °C and resuspended 1.3 ml Buffer 3 (10 mM pH8 Tris, 1 mM EDTA, 0.5 mM EGTA, 0.5% N-lauroyl-sarcosine) supplemented with protease inhibitors (PMSF, aprotinin, leupeptin, pepstatin). Suspended cells were sonicated using the Bioruptor (Diagenode) with the setting (high, 30-sec intervals for 30 min) and chromatin were collected after centrifuge at 14,000 rpm for 10 min at 4 °C. The size of reverse cross-linked DNA was 300–500 bp determined by DNA electrophoresis. Each chromatin sample was quick-frozen in liquid nitrogen and stored at −80 °C.

### Chromatin-DNA quantitation

We mixed 20 μl of chromatin with 180 μl of T_50_E_10_S_1_ (50 mM Tris pH 8.0, 10 mM EDTA, 1% SDS) at 65°C for overnight for reverse-crosslinking. We added RNase A to chromatin samples and then incubated them for 1 h at 37°C to eliminate RNA from samples. Samples were incubated with Proteinase K for 1 h at 55°C. Extract samples were mixed with phenol:chloroform:isoamyl-alcohol and sequentially added 20 μg glycogen. Purified chromatin-DNA were collected by Ethanol/NaCl precipitation and their concentration were determined with a Qubit Fluorometer (Thermo Fisher Scientific).

### Chromatin-immunoprecipitation (ChIP)

Dynal magnetic beads (Thermo Dynabeads Protein A: 10001D) were incubated with 20 μl of H3K4me3 or H3K27me3 antibodies for 8 hours at 4 °C. We prepared chromatin solution mixed with 20 or 10 μg chromatin for H3K4me3 or H3K27me3 antibodies with 5% S2 chromatin in incubation buffer (3% Triton X-100, 0.3% sodium deoxycholate, 15 mM EDTA) supplemented with protease inhibitors (PMSF, aprotinin, leupeptin, pepstatin). We combined the chromatin with antibody-coupled beads and rotated the mixture overnight at 4°C. Each bead was washed 6 times by ice-cold RIPA (50 mM Hepes-KOH pH7.5, 10 mM EDTA, 0.7 % sodium deoxycholate, 1 % NP-40, 0.5 M LiCl) supplemented with protease inhibitors (PMSF, aprotinin, leupeptin, pepstatin) and additionally washed Tris-buffer (10 mM Tris pH8.0, 1 mM EDTA, 50 mM NaCl). We eluted chromatin with 200 μl of T_50_E_10_S_1_ (50 mM Tris pH 8.0, 10 mM EDTA, 1% SDS) at 65°C for 30 min by shaking at maximum speed. Beads were centrifuged at 14,000 rpm for 1 min and placed on the magnetic stand. After revers-crosslinking described above, purified chromatin-DNA concentration was determined with a Qubit Fluorometer (Thermo Fisher Scientific).

### ChIP-Seq analysis

We mapped all precipitated read to *Bos taurus* (bosTau8) and *D. melanogaster* (dmel5.41) genome by bowtie2-2.1.0 and processed generated files by Samtools^75,76^. Normalization factors were calculated based on the number of reads mapped to the *D. melanogaster* spike-in genome (Supplementary Table 1)^34^. As we expected, around 90% reads precipitated by H3K27me3 antibody were mapped to *B. taurus* genome, while H3K4me3 were mapped relatively low percentage (32.18–63.12%). This is because H3K4me3 antibodies successfully precipitated marks of both *T. annulata* and *B. taurus* histone 3 tails and the unmapped reads to either *B. taurus* or *D. melanogaster* genomes were mapped to the *T. annulata* genome (data not shown). Peak calling was performed using Adaptive Local Threshold (ALT) equipped in Easeq (windowsize:3000 bp for H3K27me3, 1500 bp for H3K4me3, p-value:1E-5, false discovery rate (FDR):1E-5, Log2fold diff :2, merge within:100bp) or MACS2 algorithm and subsequently annotated to *B. taurus* transcripts (UMD3.1.1, n=44681)^77,78^. For clustering of each Chip-seq, we performed MACS2 peak call and clustering of all data by Bioconducter DiffBind package^79^. DiffBind clustering of peaks and high Pearson’s correlations coefficient (r >0.95) between 2 biological replicates of each macrophage and antibody indicated that the ChIP-seq had worked correctly (Supplementary Fig. 1b and c). We calculated H3K27me3 and H3K4me3 ratio of quantified value of each gene around +/- 2 kb from TSS between virulent and attenuated cell. Venn diagrams of the histone mark positive transcripts between virulent and attenuated macrophages were generated by R. All raw data of ChIP-seq and RNA-seq have been uploaded to European Nucleotide Archive (www.ebi.ac.uk/ena/) under the Study accession number PRJEB52136.

### Fibronectin binding assay

Virulent and attenuated macrophages were treated with DMSO, 1 µM UNC1999 or 1 µM UNC2400 (inactive analog) for 10 days^37,38^. We seeded treated macrophages onto fibronectin-coated plates and incubated them for 1 h at 37 °C. Bound cells after PBS wash were stained by Violet blue and lysed by lysis buffer containing Triton X-100. The number of cells was quantified by measuring absorbance at 595 nm.

### Dissemination of *T. annulata*-transformed macrophages in immune-deficient mice

*T. annulata*-transformed virulent and attenuated macrophage lines were treated *in vitro* for ten days with DMSO or 1 µM UNC1999, as described above. The macrophages (10^6^ in 200 µl PBS) were then injected subcutaneously into right flank region of ‘alymphoid’ Rag2/γC double-deficient (T-, B-, NK-) mice and surveyed for 3 weeks^35^. Four groups each containing five age and sex matched mice were defined: two treatment groups including virulent/attenuated Ode treated with EZH2 inhibitor and two control groups including virulent/attenuated Ode treated only with drug vehicle (solvent). During the three-week study period, mice in the treatment groups were orally administered with UNC1999 (100 mg/kg/day) dissolved in a solution of Tween® 80 (Sigma-Aldrich, Germany) and carboxymethyl cellulose sodium salt (Sigma life science, USA) to facilitate oral gavage. The control mice received 100 µl of solvent per day. At the end of study, mice were humanely sacrificed and organs were dissected and kept in PBS at −20°C.

### Absolute quantification of *T. annulata-*infected cells in mice tissues

A single copy *T. annulata* gene (apical merozoite antigen 1, TA02980) was cloned into pJET 1.2/blunt cloning vector using CloneJET PCR Cloning Kit (Thermo Fisher Scientific). To measure the load of *T. annulata-*infected macrophages in each tissue, quantitative PCR technology was applied to genomic DNA extracted from mouse tissue (QIAmp DNA mini kit). Quantification were done based on the methodology described elsewhere^80^. Briefly, a standard curve was generated from crossing points (Cp) measured from serial dilutions of pJET-ama plasmid construct with known plasmid copy numbers using qPCR assay. The Cps from 1 ng DNA of each tissue sample were used to estimate parasite gene copy numbers based on the equation derived from plotted data of the standard curve graph.

### Library preparation and sequencing

We maintained virulent and attenuated macrophages with 1 μM PRC2 inhibitor for 10 days, after which a total of 2.5 × 10^6^ cells were collected for RNA extraction using mirVana miRNA isolation Kit (Thermo Fisher Scientific, catalogue number AM1560) according to the manufacturer protocol. Strand-specific RNA-sequencing (ssRNA-seq) libraries were prepared using the Illumina Truseq Stranded mRNA Sample Preparation Kit (Illumina, catalogue number RS-122-2101) following the manufacturer’s instructions. Briefly, 1 ug of total RNA was used to purify mRNA using poly-T oligo-attached magnetic beads. mRNA was then fragmented and cDNA was synthesized using SuperScript III reverse transcriptase (Thermo Fisher Scientific, catalogue number 18080044), followed by adenylation on the 3’ end, barcoding and adapter ligation. The adapter ligated cDNA fragments were then enriched and cleaned with Agencourt Ampure XP beads (Agencourt, catalogue number A63880). Libraries validation was conducted using the 1000 DNA kit on 2100 Bioanalyzer (Agilent Technologies, catalogue number 5067-1504) and quantified using Qubit (Thermo Fisher Scientific, catalogue number Q32850). ssRNA libraries were sequenced on Illumina Hiseq4000.

### RNA seq analysis

All reads were trimmed using Trimmomatic and then aligned to the *Bos taurus* UMD.3.1 genome with HISAT2^81^. BAM files converted by Samtools, as described above and were processed by HTSeq-count tools^82^ to quantify mapped reads on each gene. We obtained a DEGs lists among each group with logFC > ±1 and padj < 0.05 using separately EdgeR and Limma of NetworkAnalyst or TCC-GUI^83,84^ and made intersections of these two lists to identify more stringent DEGs. Treemap of Gene ontology analysis were visualized by Revigo^85^.

### Gene expression measurements by qRT-PCR

Total RNA was extracted from cells using RNAeasy® plus mini kit (QIAGEN). Complementary DNA (cDNA) synthesis was done by using 1 µg RNA as template and the GoScript™ reverse transcriptase kit (Promega, Cat. No. A5001, USA), in 20 µl reaction volume, following the manufacturer’s instructions. To perform qRT-PCR target sequences were amplified in a SYBR green PCR master mix (ThermoFisher Scientific, Cat. No. 4309155) mixed with diluted cDNA (1:20), double distilled water and primers. The PCR was run in LightCycler® 480 instrument (Roche) and results analysed in Microsoft Excel. Finally, the 2^−ΔΔCT^ methodology was employed to estimate relative gene expression levels^86^. The expression of Bovine GAPDH was used as the internal control for normalization. Graphs were prepared in GraphPad Prism V 8.4.0.

### Ethics Statement

A detailed protocol (number 12-26) describing the above mice experiments was first submitted to and approved (number CEEA34.GL.03312) by the ethics committee for animal experimentation at the University of Paris-Descartes. The university ethics committee is registered with the French National Ethics Committee for Animal Experimentation that itself is registered with the European Ethics Committee for Animal Experimentation. The right to perform the mice experiments was obtained from the French National Service for the Protection of Animal Health and satisfied the animal welfare conditions defined by laws (R214-87 to R214-122 and R215-10) and GL was responsible for all experiment as he holds the French National Animal Experimentation permit with the authorization number (B-75-1249). All experimental procedures in KAUST were approved by Institutional Biosafety and Bioethics Committee (IBEC) in KAUST (IBEC number: 22IBEC029).

## Author Contributions

**TS** performed all wet-bench experiments, ChIP-seq bioinformatic analyses and wrote the manuscript with editorial inputs from **AP**, **MW** and **GL**. **ST** performed all mouse experiments including estimation of tumor dissemination and qRT-PCR analysis. He also did the wet-bench experiments requested by the reveowers. **ZR** made the RNA-seq libraries that were bioinformatically analyzed by **TS**, **HRA**, **AK** and **TM**. The study was conceived by **MW** and **GL** and supported by a joint grant awarded to **AP** and **GL**.

## Acknowledgements

**TS** and **ST** acknowledge a LabEx ParaFrap postdoctoral fellowship and **GL** acknowledges ANR-11-LABX-0024 support and core funding from INSERM and the CNRS. **GL** and **AP** acknowledge financial support received from KAUST in form of a joint (OCRF-20146CRG4) grant and **AP** a BAS/1/1020-01-01 grant. We thank Raphael Margueron at the Curie Institute for his assistance with the ChIP-seq experiments; the Next Generation Sequencing Platform at the Curie Institute for ChIP sequencing, and members of the KAUST Bioscience Core Laboratory (BCL) for generating the raw RNA-seq datasets. We also thank Axel Martinelli at Hokkaido University for instruction on RNA-seq analysis and Mads Lerdrup at the University of Copenhagen for guidance on ChIP-seq analysis by EaSeq.

---

**Supplementary Figure 1.**
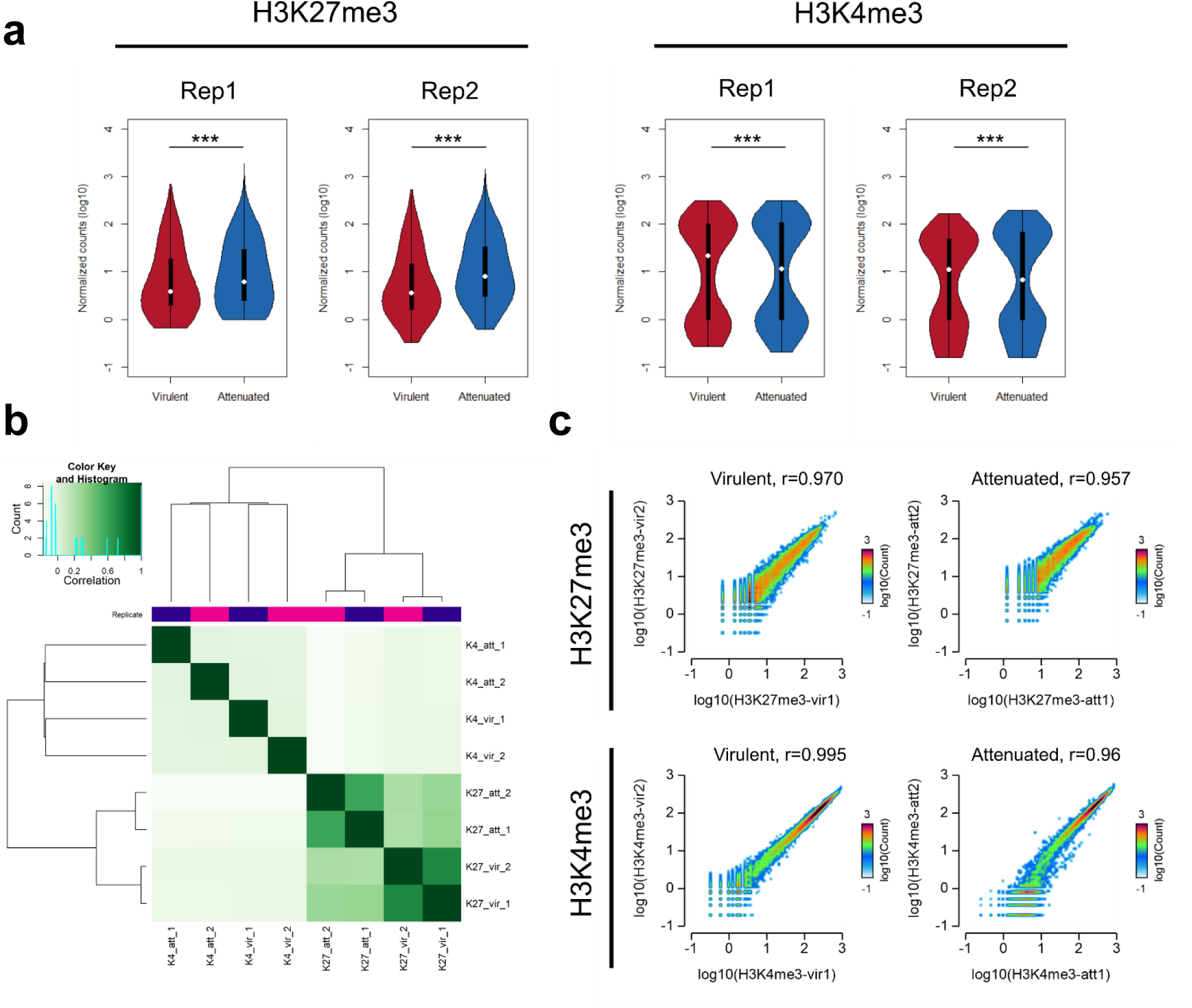
Clustering of each ChIP-seq reads and scatter plot. **a)** Violin plots represent the read counts on TSS around +/- 2kb of each gene obtained from H3K27me3 and H3K4me3 ChIP-seq. ***, Wilcoxon rank-sum test p-value < 2.2e-16. **b)** Clustering of each ChIP-seq samples visualized by Bioconductor DiffBind package. **c)** 2D density plots comparing H3K27me3 and H3K4me3 between ChIP-seq biological replicates. Read counts for each gene in replicate 1 (on X-axis) and replicate 2 (on Y-axis) are plotted. The r values represent the Pearson correlation coefficients.

**Supplementary Figure 2.**
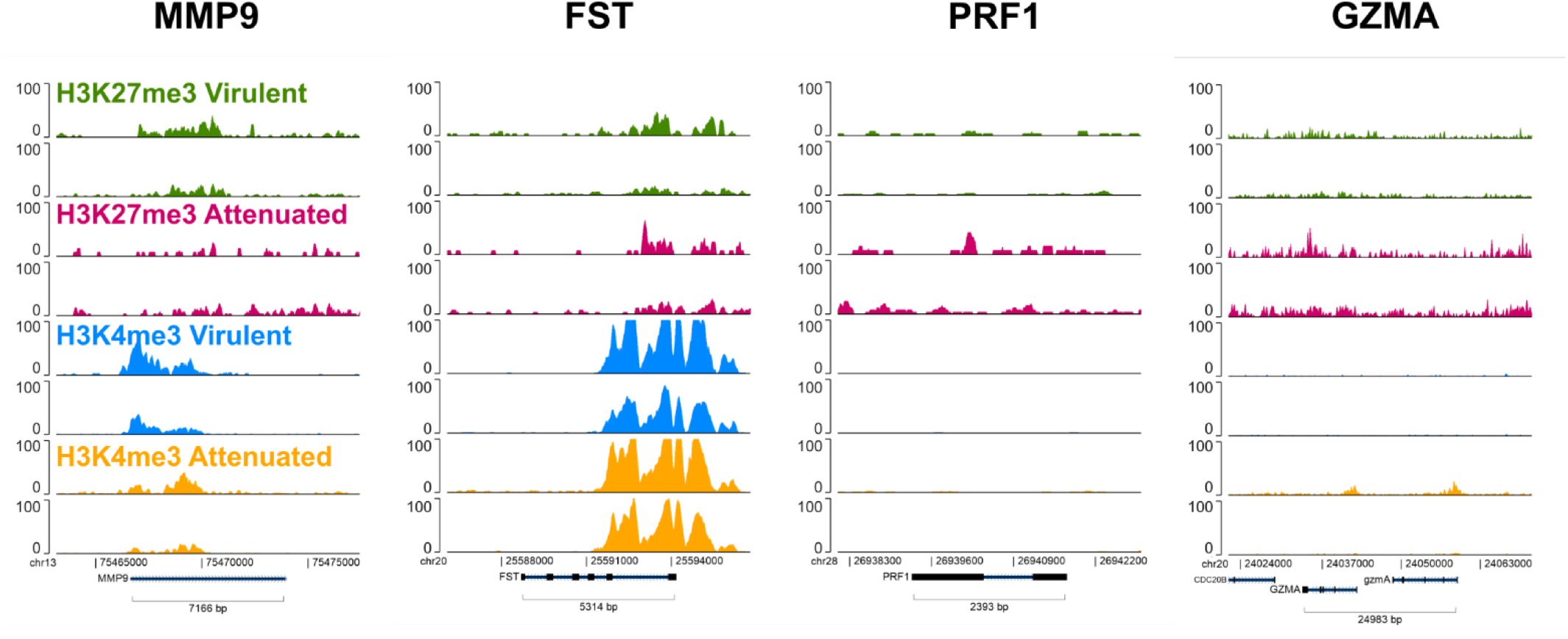
Genomic distribution of H3K27me3 and H3K4me3 marks around 4 representative genes of the 21 common upregulated genes relative to Figure 5. Profile of each mark on the *MMP9*, *FST*, *PRF1*, *GZMA* genes.

**Supplementary Figure 3.**
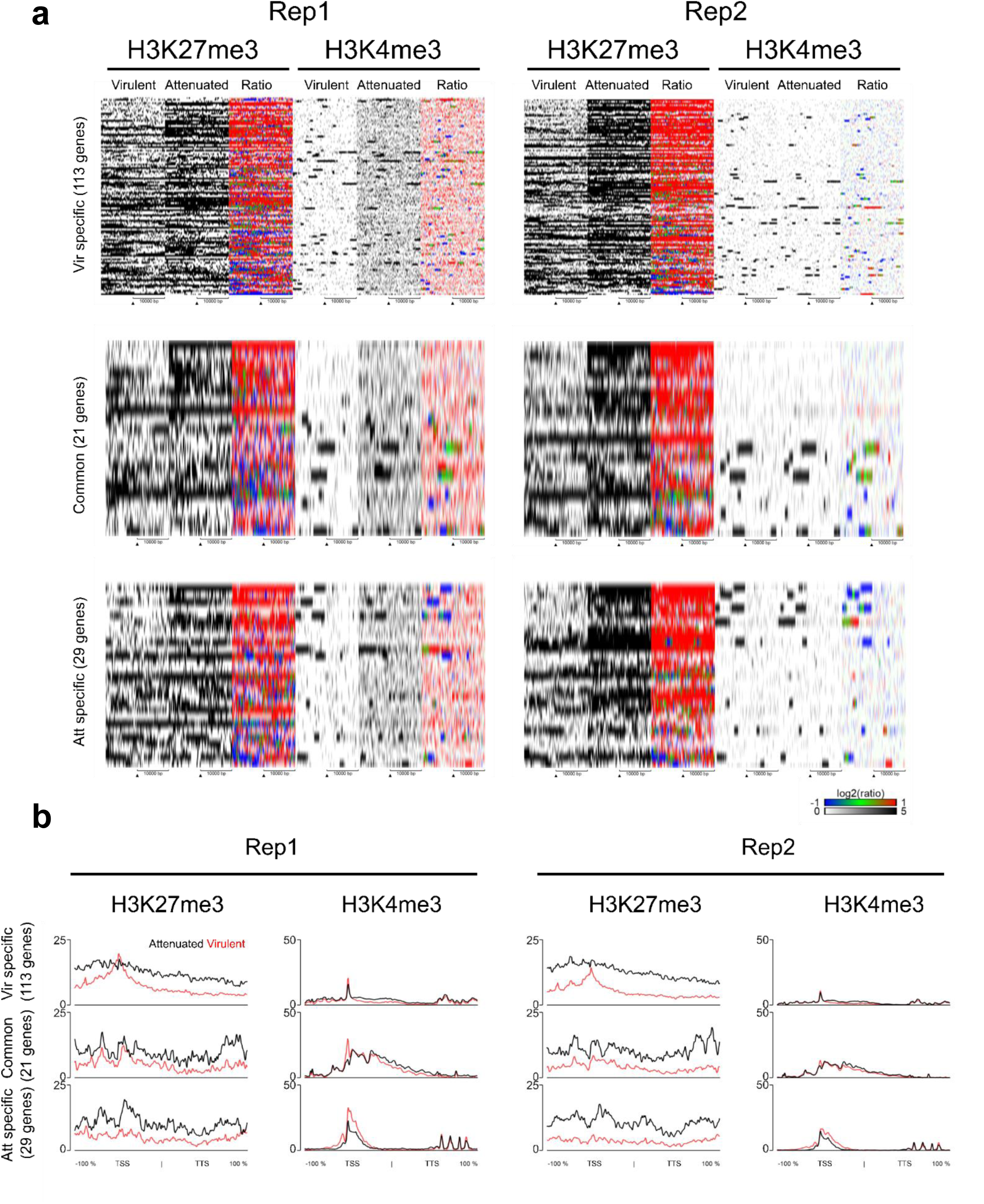
H3K27me3 and H3K4me3 profiles of upregulated genes upon PRC2 inhibitor related to Fig. 5b. **a)** H3K27me3 and H3K4me3 heat maps around TSS +/- 10 kb of 3 groups of genes related to Fig.5b in virulent or attenuated macrophages. Genes were sorted by the ratio of H3K27me3 signal log2 (Att/Vir). **b)** Average H3K27me3 and H3K4me3 signal intensities along the 3 groups of genes defined in Fig.5b in virulent or attenuated macrophages. The X-axis represents a window centered on each gene, spanning from the TSS to the TTS with an additional ±100% flanking region. Black lines represent signals from virulent macrophages, whereas red lines represent those from attenuated macrophages.

**Supplementary Figure 4.**
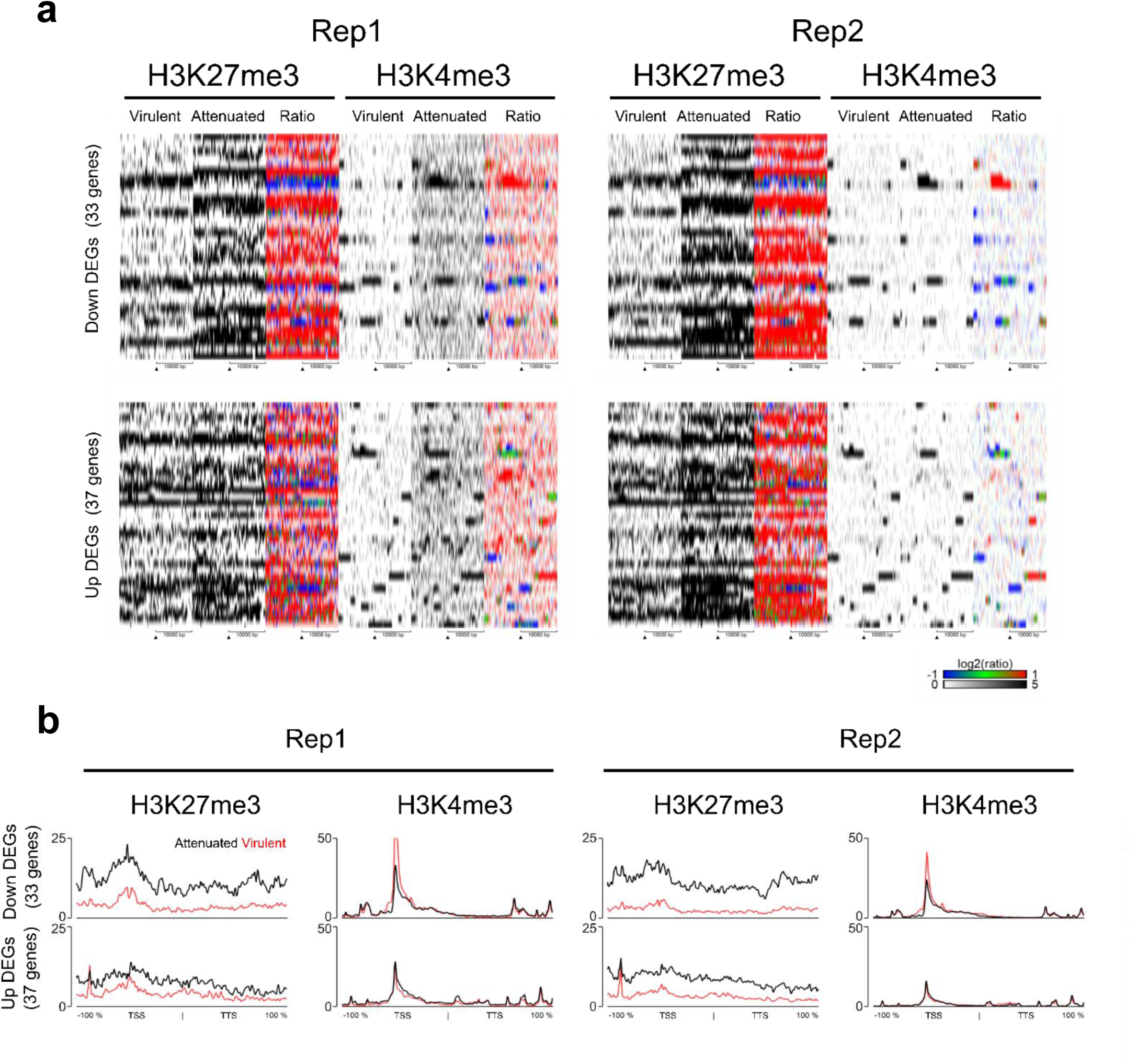
H3K27me3 and H3K4me3 profiles of Down and Up DEGs in attenuated macrophages. **a)** H3K27me3 and H3K4me3 heat maps around TSS +/- 10 kb of Down and Up DEGs in attenuated macrophages. **b)** Average H3K27me3 and H3K4me3 signal intensities along genes that are significantly up- or downregulated in attenuated macrophages. The X-axis represents a window centered on each gene, spanning from the TSS to the TTS with an additional ±100% flanking region. Black lines represent signals from virulent macrophages, whereas red lines represent those from attenuated macrophages.

**Supplementary Figure 5.**
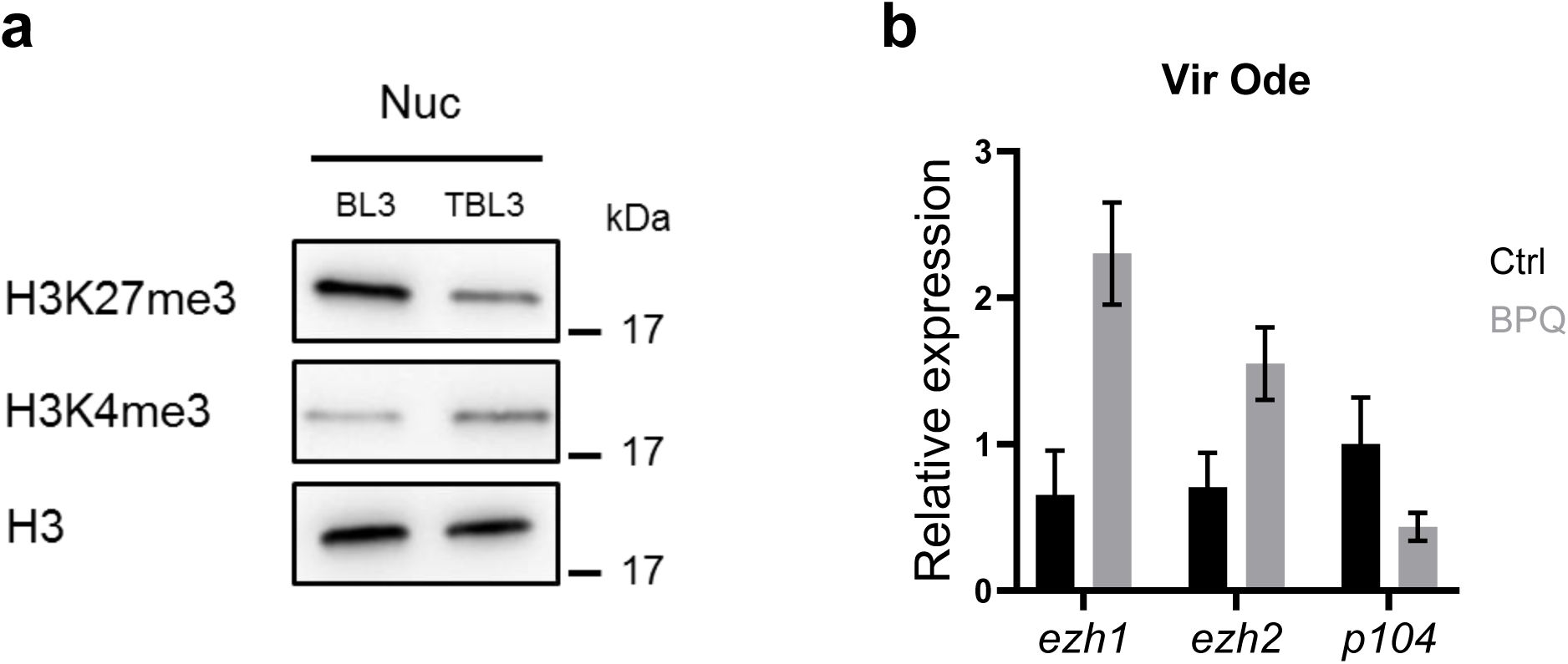
*Theileria annulata* infection downregulates PRC2 complex function. **a)** Western blot analysis of nuclear extracts from uninfected (BL3) and infected (TBL3) bovine B cells shows a global decrease in H3K27me3 levels in TBL3. **b)** Elimination of the parasite by treating *T. annulata*-transformed macrophages (TaC12) with buparvaquone (BPQ) results in increased *ezh1* and *ezh2* mRNA levels, supporting the data shown in panel (a). Expression of the *p104* gene encoding a major *T. annulata* schizont surface protein was also measured to demonstrate the effect of BPQ on parasite clearance.

**Supplementary Table 1. Summary of ChIP-seq experiments**

**Supplementary Table 2. List of differentially expressed genes corresponding to Fig 5b**

**Supplementary Table 3. List of primers for qRT-PCR**

**Supplementary Table 4. List of GO terms of DEGs corresponding to Fig 5c**

